## Supplemental Figures for "Matrix viscoelasticity regulates dermal fibroblast activation in a three-dimensional fibrillar microenvironment"

### Supplementary Figures

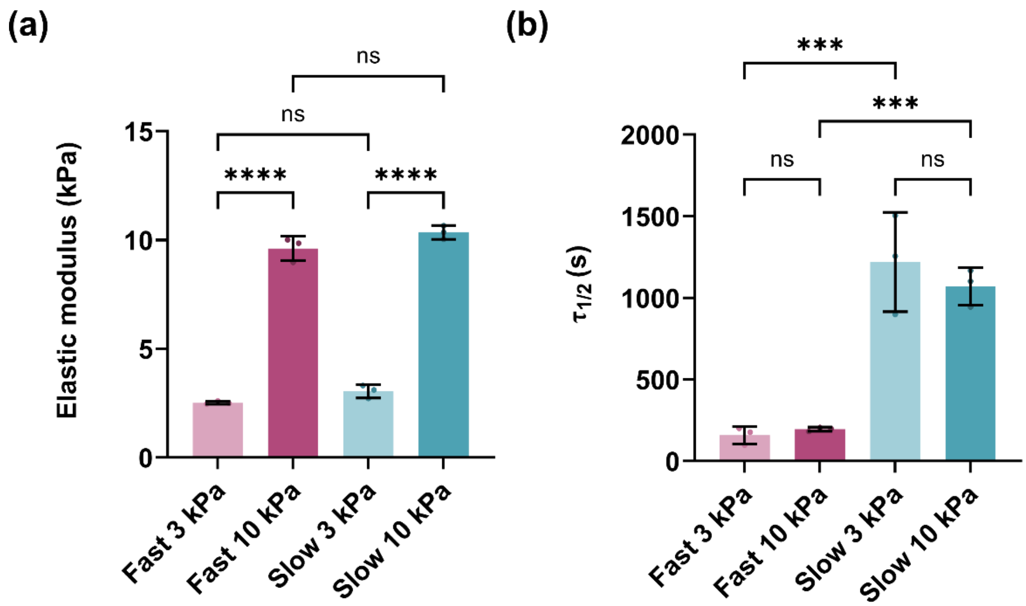


**Supplemental Figure 1: RGD-alginate matrices exhibit independently tunable elastic modulus and stress relaxation rates.** (a) Elastic modulus measurements of RGD-modified alginate hydrogels. (b) Stress relaxation rates of RGD-modified alginate hydrogels, represented by $\tau_{1/2}$. Data presented using mean ± standard deviation for n = 3 replicates per condition and analyzed using one-way ANOVA, followed by Šidák's multiple comparison test. A p-value of less than 0.05 was considered statistically significant. *** $p < 0.001$, **** $p < 0.0001$, ns; not significant.

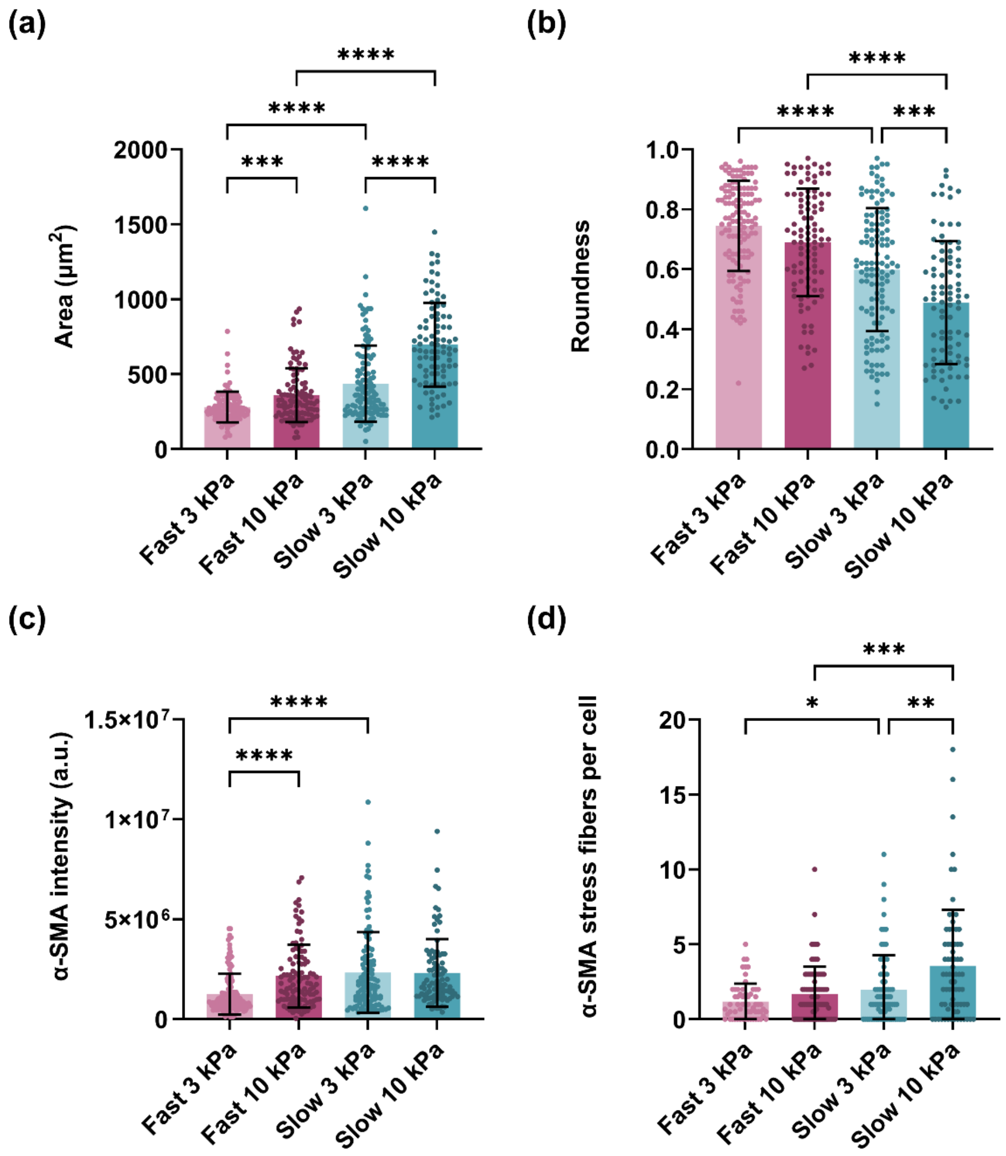


**Supplemental Figure 2: Matrix stiffness and stress relaxation regulates fibroblast morphology and cytosolic α-SMA expression in RGD-alginate matrices.** (a) Quantifications of fibroblast area. Fibroblast area increases in matrices in both a stiffness-dependent and stress relaxation-dependent fashion. (b) Quantifications of fibroblast roundness. (c) Quantifications of cytosolic α-SMA expression. Cytosolic α-SMA was upregulated with increased elastic modulus and decreased stress relaxation rate. (d) Quantifications of α-SMA stress fibers. Data presented using mean ± standard deviation for $n = 3$ biological replicates per matrix condition. Statistical significance was determined by ANOVA, followed by Dunnett's multiple testing correction, with p-values < 0.05 considered to be statistically significant. * $p < 0.05$; ** $p < 0.01$; *** $p < 0.001$; **** $p < 0.0001$. If there is no statistical significance indicator bar, comparisons were not statistically significant ($p > 0.05$).

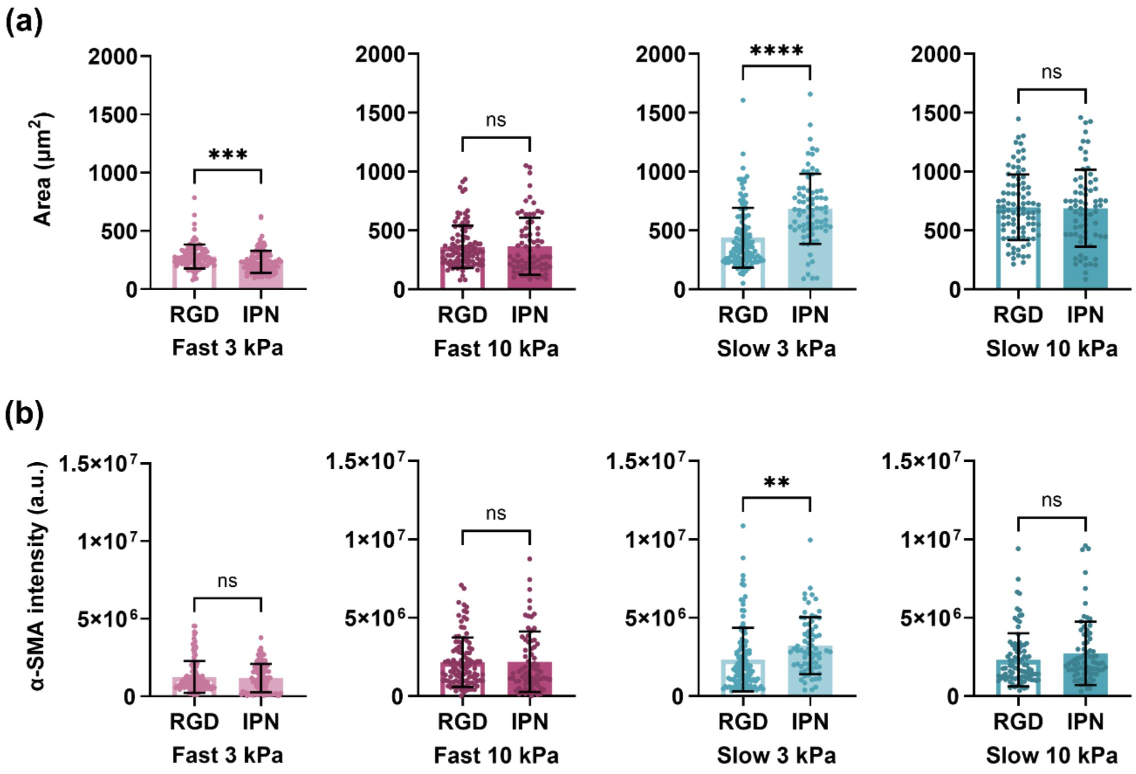


**Supplemental Figure 3: Fibroblast area and α-smooth muscle actin expression are comparable in RGD-alginate matrices and alginate-collagen I interpenetrating networks.** (a) Comparisons of cell spread area between RGD-alginate matrices and IPNs. (b) Comparisons of α-SMA expression measured in RGD-alginate matrices and IPNs. Data presented using mean ± standard deviation for n = 3 biological replicates per matrix condition and analyzed using unpaired t-tests followed by Welch's correction. A p-value of 0.05 or less was considered statistically significant. ** $p < 0.01$; **** $p < 0.0001$; ns, not significant.
